# Retinoid-impregnated nanoparticles enable control of bone growth by site-specific modulation of endochondral ossification in mice

**DOI:** 10.1101/2024.11.08.622655

**Authors:** Masatake Matsuoka, Kenta Uchibe, Ningfeng Tang, Hongying Tian, Akiko Suzuki, Takeshi Oichi, Yu Usami, Ivan Alferiev, Satoru Otsuru, Joshua M. Abzug, John E. Herzenberg, Maurizio Pacifici, Motomi Enomoto-Iwamoto, Michael Chorny, Masahiro Iwamoto

## Abstract

Growth-plate (GP) injures in limbs and other sites can impair GP function and cause deceleration of bone growth, leading to progressive bone lengthening imbalance, deformities and/or physical discomfort, decreased motion and pain. At present, surgical interventions are the only means available to correct these conditions by suppressing the GP activity in the unaffected limb and/or other bones in the ipsilateral region. Here, we aimed to develop a pharmacologic treatment of GP growth imbalance that involves local application of nanoparticles-based controlled release of a selective retinoic acid nuclear receptor gamma (RARγ) agonist drug. When RARγ agonist-loaded nanoparticles were implanted near the medial and lateral sides of proximal tibial growth plate in juvenile C57BL/6j mice, the GP underwent involution and closure. Overall tibia length was shortened compared to the contralateral element implanted with drug-free control nanoparticles. Importantly, when the RARγ agonist nanoparticles were implanted on the lateral side only, the adjacent epiphysis tilted toward the lateral site, leading to apical angulation of the tibia. In contrast to the local selectivity of these responses, systemic administration of RARγ agonists led to GP closure at many sites, inhibiting skeletal growth over time. Agonists for RARα and RARβ elicited no obvious responses over parallel regimens. Our findings provide novel evidence that RARγ agonist-loaded nanoparticles can control activity, function and directionality of a targeted GP, offering a potential and clinically-relevant alternative or supplementation to surgical correction of limb length discrepancy and angular deformities.

**Lay summary:** Growth-plates (physes), which are cartilage tissues near the ends of bones, support normal bone growth in children. Growth plate injures in limbs and other sites can impair growth plate function, leading to inhibited or imbalanced bone growth, skeletal deformities, decreased motion, discomfort or pain. At present, surgical interventions are the only means available to correct these conditions. Here, we aimed to develop a pharmacologic treatment for bone growth imbalance. Nanoparticles loaded with a selective agonist for the retinoic acid nuclear receptor gamma were prepared and implanted near the tibial growth plate in juvenile mice. The growth plate underwent involution and closure, and overall tibia length was shortened compared to the contralateral element implanted with drug-free control nanoparticles. Importantly, when the same drug nanoparticles were implanted in only one side of the tibia, the tibia was tilted toward the injection site. Our findings provide novel evidence that retinoic acid receptor gamma agonist-loaded nanoparticles can control activity, function and directionality of a targeted growth plate, offering a potential and clinically-relevant alternative or supplementation to surgical correction of limb length imbalances and deformities.

## Introduction

Growth plates (physes) are located at both ends of long bones and are essential for bone growth.^1,2^ Growth plate (GP) damage by fractures, tumors and other conditions or injuries can result in deceleration of skeletal lengthening and decrease in bone formation.^3,4^ On the other hand, diaphyseal fractures in the femur or tibia often stimulate excessive bone growth,^5,6^ presumably due to regional activation of the GP function. Importantly, both conditions can trigger imbalances in length and shape compared to unaffected elements or among neighboring bones, leading to progressive deformity of the skeleton and significant physical problems.^7–9^ At present, surgery is the only practical means to locally correct deformities due to impaired bone growth.^7,9^ Such surgical procedures are invasive and impose a substantial burden to children and their families, necessitating the development of alternative and non-surgical therapeutic strategies.

Within the GP, chondrocytes undergo a series of functional changes driving bone formation, a process called endochondral ossification. These chondrocytes actively proliferate, produce abundant cartilage matrix, further differentiate into hypertrophic chondrocytes, and induce matrix degradation and calcification.^2,10^ The GP is then invaded by newly formed blood vessels and osteoclasts and is eventually replaced by bone. The endochondral ossification process is controlled by multiple factors including systemic and local soluble factors (vitamins, hormones, growth factors and cytokines), extracellular matrix molecules, nutrition and mechanical loading.^2,10,11^

The active form of vitamin A, all-*trans*-retinoic acid, exerts a key role in numerous biological processes including skeletogenesis and myogenesis.^12–16^ Its action is mediated by various combinations of nuclear hormone receptors that primarily act as transcription factors.^17^

Heterodimers composed of a RAR (retinoic acid receptor) and RXR (retinoic acid receptor X) subunits translocate into the nuclei, bind to specific DNA sequences featuring a retinoic acid responsive element (RARE) and regulate target gene expression (genomic action).^13,18,19^ RAR signaling and function in different tissues and developmental stages are governed by the specific expression patterns of RARα, RARβ and RARγ and the availability of intracellular active retinoids.^13,19^ Previous findings have demonstrated that RARγ is a dominant RAR isoform in growth plate cartilage and supports matrix production as an unliganded transcriptional repressor.^14^ On the other hand, excessive activation of RARγ inhibits chondrogenesis and ectopic endochondral ossification, and this action has been studied preclinically and clinically to prevent acquired and genetic heterotopic ossification.^20–24^ In comparison, pan-antagonists of RARs potentiate chondrogenesis and bone defect healing^25^ and are able to counteract hedgehog inhibitor-induced GP closure.^26^ These findings led us to hypothesize that local pharmacologic modulation of RARγ function could provide a therapeutic modality for controlling bone growth via modulation of GP activity.

Targeted drug delivery using nanocarriers has been shown to improve efficacy and minimize side effects on non-target tissues.^27^ Nanocarrier-based formulations can deliver drugs to their local sites of action while protecting labile compounds from premature degradation, increasing bioavailability, limiting peripheral dissemination and addressing solubility issues. In this study, using the mouse model, one of the most commonly used animal models to study skeletal growth, we tested polymeric nanoparticles (NP) as biodegradable drug carriers to generate proof-of-concept evidence that this therapeutic approach could be effective against GP activity imbalances. We also examined the molecular actions of selective agonists of RARγ on GP activity and evaluated NP-based drug delivery as an approach to control and modulate targeted bone growth. Our results provide novel and far-reaching evidence that nanoparticles delivering selective RARγ agonists to a growing skeletal element can restrict bone lengthening and modulate growth directionality, providing an effective and seemingly safe tool to treat growth imbalances and deformations on their own and/or in conjunction with surgery.

## Materials and methods

### Animals

All animal experiments were performed following institutional guidelines and were approved by the Institutional Animal Care and Use Committees at Children’s Hospital of Philadelphia (959) and University of Maryland, Baltimore (0317003 and 0120005). Animals were housed in a ULAR supervised animal facility with a 12-hour light/dark cycle in a temperature (22±1°C) and humidity (55±5%) controlled room. Animals were provided hygienic animal bedding; all cages contained wood shavings, bedding and cotton pads. The health status of each animal was monitored throughout the experiments by investigators as well as by animal veterinary technicians and veterinarians as per the institutional guidelines. The mice were free of all viral, bacterial, and parasitic pathogens during the experimental schedule. Body weight and activity of the mice indicating their general health were monitored before and throughout the experiments. RARγ deficient mice^28^ were kindly provided by Dr. Pierre Chambon and Norbert B. Ghyselinck (INSERM, France). Genotyping procedure was described previously.^14^ C57BL/6J mice (#000664) and Ai6 mice (Cre-sensitive ZsGreen reporter mice, B6.Cg-Gt(ROSA)26Sortm6(CAG-ZsGreen1)Hze/J, #007906) were purchased from the Jackson Laboratory (Bar Harbor, ME). Mice were randomly divided and allocated to control and drug groups. Drug response was assessed by bone length differences (primary outcome) and histological changes (secondary outcome). Animal treatment and sample analysis were performed in an unblinded manner.

### Preparation and systemic administration of retinoids

NRX 204647^20^ (RARγ agonists, CAS1351452-80-6) were synthesized by Atomax Chemicals (Shenzhen, China). Retinoic acid, BMS453 (RARβ agonist, CAS166977-43-1) and CD2665 (RARβ/γ antagonist, CAS 170355-78-9) were purchased from Tocris Biosciences (Bristol, UK). AGN195183 (RARα agonist, CAS367273-07-2) was obtained from NuRx Pharmaceuticals (Irvine, CA). RARγ antagonist 7C was synthesized by Atomax Chemicals (Shenzhen, China) according to the world patent application WO 2005/066115 A2. The concentrations of retinoids used for animal experiments were 1 mg/kg for NRX 204647 and 4 mg/kg for all other compounds unless indicated. Stock solutions of retinoids in DMSO (D2650; Sigma-Aldrich, St. Louis, MO) were stored at −30°C under argon. Before administration, aliquots of stock solution were mixed with three-fold volume of corn oil (C8267; Sigma-Aldrich, St. Louis, MO) for each dose, and administered to mice. Vehicle control mice received 10 μl DMSO plus 30 μl corn oil in the same manner. C57BL/6J mice were treated with vehicle or retinoids at 11, 13 days of age by an intraperitoneal injection and 15 and 17 days of age by oral gavage using a 20-gauge gavage needle (Fine Science Tools, Foster City, CA) at indicated time points. The mice were euthanized 1, 3, 5 or 8 days after the treatment started. The body length was determined from nose to base of the tail by a caliper. The hind limbs were harvested and subjected to histological assessments.

C57BL/6J female and male mice were also treated with vehicle or NRX204647 every other day starting at 14 days of age by an intraperitoneal injection, the tibia length was radiologically measured 8 days after the drug treatment started. Some mice received a subcutaneous injection of 50 mg/kg 5-ethynyl-2’-deoxyuridine (EdU) (Thermo Fisher Scientific, Waltham, MA) 2 hours before euthanization. The tibia and femur lengths were determined by measuring a linear distance between the proximal and distal ends of the articular cartilage.

### Preparation and local administration of NRX204647

Based on the chemical structure and receptor binding selectivity, NRX204647 was chosen as RARγ agonist to be administered in PLA-based NPs (NRX-NP). NPs were formulated by a modification of the emulsification-solvent evaporation method using serum albumin as a colloidal stabilizer.^29^ In brief, NRX204647 (15 mg) was co-dissolved with 200 mg of PLA (Mn = 19 kDa, PDI = 1.8, Birmingham Labs, Birmingham, AL) in chloroform (5 ml). The organic phase was emulsified by sonication in an aqueous solution containing 150 mg of human serum albumin (Octapharma AB, Stockholm, Sweden) dissolved in 10 ml of deionized water, and the organic solvent was removed by rotary evaporation under reduced pressure. NPs were passed through a sterile 5.0-µm membrane, lyophilized with glucose (5% w/v) as a cryoprotectant, stored at -80°C, and reconstituted in deionized water before use. The drug loading of NP was determined spectrophotometrically following extraction in *sec*-butanol against a suitable calibration curve. NP size measurements were performed by dynamic light scattering.

Fluorescently labeled NP were formulated as above by incorporating a PLA-BODIPY_558/568_ conjugate as described elsewhere^30^. A similar process was used to make drug-free (blank) NP formulations or NP loaded with 4-hydroxy-tamoxifen (4OH-Tx, Sigma-Aldrich Inc., St. Louis MO). The NPs were diluted with sterilized saline making at the desired concentration: NRX-NP, 0.8 μg/μl; 4OH-Tx-NP, 5μg/μl).

The incision was made to expose the left proximal epiphysis of of the tibia of 3-week-old C57BL/6j or Ai6 female mice, and NRX-NP (1.6 μg retinoid/2 μl/site), 4-OH-tamoxifen (10 μg/2μl/site)-loaded NPs, or NRX204647 solution (1.6 μg/2ul DMSO/site), respectively were weekly injected in the vicinity of the GP of the left proximal tibiae followed by suture of the incision. Right tibiae received empty NPs (Blank-NP) or vehicle solution in the same manner. The NPs were injected from the medial site only or from both medial and lateral sites per time. The tibias were analyzed at 5 weeks of age. The BODIPY-NPs were injected in the vicinity or into the medullary cavity of the tibia and the tissues were harvested 1 or 7 days after injection.

### Histological and immunohistochemical analyses

Tissue samples were fixed in 4% paraformaldehyde overnight at 4°C, decalcified in 10% EDTA or 10% formic acid processed for paraffin embedding, and serially sectioned at 6 μm-thickness. Sections were stained with hematoxylin and eosin (HE) for general profiling. The tibiae that received the BODIPY-NPs were fixed in 4% paraformaldehyde overnight at 4°C and embedded in optical cutting temperature compound for frozen sectioning. Undecalcified sections were made and by Kawamoto’s method,^31^ and the distribution of BODYPI-NPs was examined. For immunohistochemical staining, the sections were baked for 1 h at 60°C and then de-paraffinized. Sections were incubated with 400 IU/mL hyaluronidase (VDIPEN staining) or 0.1% trypsin (collagen 10 staining) for 30 min at 37°C, or treated with 10 mM sodium citrate (pH 6.0) for 10 min at 90 °C (Tgm2 staining) with blocking solution (0.1M NaPB, 1% BSA and 10% normal goat serum) for 1 h at 22°C, and then with primary antibodies against aggrecan degradation products (VDIPEN, 1:3000; provided by Dr. J. Mort, Shriners Hospital, Canada),collagen 10 (1:1000, ab58632; Abcam, Cambridge, MA) or Tgm2 (1:300, #15100-1-AP; Proteintech Group, Inc, Rosemont, IL) overnight at 4°C. The antibodies were visualized using rabbit specific HRP/DAB (ABC) detection kit (ab64261; Abcam). After detection, sections were counter stained with 0.5% methyl green. The results were observed and captured using a BZX700 microscope (Keyence, Itasca, IL, USA).

### RNAscope

The hind limbs were fixed with 2% formaldehyde and dehydrated by increasing sucrose percentage to 30% and then embedded in the optimal cutting temperature (O.C.T.) compound (Thermo Fisher Scientific). Undecalcified frozen sections were mounted on the Kawamoto’s film and subjected to in situ hybridization using the RNAscope® Fluorescent Multiplex Kit (Advance Cell Diagnostic Inc. Newark. CA). We used predesigned probes (*Cyp26b1#454241*, *Tgm2 #537801*, *Ihh #413091* and *Adamts9 #400441*), obtained from the Advance Cell Diagnostic Inc. company. The results were observed and captured using a BZX700 microscope (Keyence). The images were processed using ImageJ2 version 2.14.

### Chondrocyte culture and the in vitro assays

Primary mouse chondrocytes were isolated from 3-5-day-old C57BL/6J wild type mice and homozygous and heterozygous RARγ null mice. The isolated cells were plated at the density of 1× 10^5^/cm^2^ in the 12 well plate and treated with ethanol, retinoic acid (100 nM), RARα agonist (AGN195183, 300 nM), RARβ agonist (BMS453, 300 nM) and RARγ agonist (NRX204647, 100 nM) for 2 days. The phase contract images of the cultures were then captured under the BZX-700 microscope (Keyence). Total RNAs from cell cultures were purified using RNeasy mini kit (Qiagen, Germantown, MD) and subjected to RT-qPCR using SYBR green (Applied Biosystems, Foster City, CA) as previously described.^32^ Average threshold cycle value (Ct value) was calculated from quadruplicate reactions. Standard curves were generated using 10-fold serial dilutions of cDNA of each gene with a correlation coefficient of >0.98. Relative expression levels were calculated based on a standard curve and normalized to *glyceraldehyde 3-phosphate dehydrogenase* (*Gapdh*). Primers are designed to amplify sequences common among known splice variants of each gene. Primer sequences used for real-time PCR amplification were as follows: 5’-AGT TTG ACC AAT AVT TGC CAC CCA AC-3’ and 5’-TCC GTC TTG ATG TGC GTT CGC T-3’ for mouse *Sox9* (NM_011448), 5′-TCT GGA AAT GAC AAC CCC AAG CAC A-3′ and 5TGG CGG TAA CAG TGA CCC TGG AAC T-3′ for 5463-5939 of mouse *Acan* (NM_007424), 5′-CTT GTG GAC AAT CCT CAG GTT TCT GTT C-3′ and 5′-TCG GTC ACC ATC AAT GCC ATC TAT G-3′ for 822-1040 of mouse *Col9a1* (NM_007740), 5′-TGC TGC TAA TGT TCT TGA CCC TGG TTC-3′and 5′-ATG CCT TGT TCT CCT CTT ACT GGA ATC C-3′ for 717-876 of mouse *col10a1* (NM_009925), 5-TCA GTC TCT TCA CCT CTT TTG GGA ATC C-3’ for 799–1160 of mouse *Mmp-13* (NM_008607.2), 5-TGTACTTTCGAGCGCAGATG-3’ and 5’-AGGCTTGTTTCATCCTCCTG-3’ for mouse *Tnfsf11* ( NM_011613.3) and 5′-AAG CCC ATC ACC ATC TTC CAG GAG-3′, 5′-ATG AGC CCT TCC ACA ATG CCA AAG-3′ for 258–568 of *glyceraldehyde 3-phosphate dehydrogenase* (NM_008084).

The freshly isolated chondrocytes were plated at the density of 1.2 x 10^5^/cm^2^ and reverse transfected on the PCR array plate (RT² Profiler™ PCR Array Mouse Extracellular Matrix & Adhesion Molecules, Qiagen). On the next day, the cultures were treated with RARα (AGN195183, 300 nM) or RARγ agonists (NRX204647, 100 nM) for 48 hrs and subjected to PCR analysis following the manufacturer’s protocol. Average threshold cycle value (Ct value) was calculated from 4 wells and normalized to that of housekeeping gene GAPDH. The experiments were repeated twice independently.

### In vitro bioassay of the NRX-NPs

The NRX-NPs were incubated in the chondrocyte culture medium (high glucose DMEM containing 10% FBS) for 2 hrs at 37°C and collected by centrifugation at 100,000 rpm for 15 min at 4°C. The supernatant was stored at 4°C until use. The precipitated NRX-NPs were suspended in the culture medium again and further incubated. This step was sequentially repeated to have 12-, 24-, 48-, 72-, 144- and 192-hrs incubation NRX-NP supernatants. The HEK293 cells that stably integrated contains a firefly luciferase gene under the control of retinoic acid response elements along with full length human *RARG* gene (BPS Bioscience, San Diego CA) were treated with a serially diluted NRX-NPs or NRX204647, the NRX-NP-supernatants or the precipitated NRX-NPs after 192hrs incubation. The luciferase activity was measured 48 hrs after treatment using the luciferase assay kit (Promega, Madison, WI).

### Gene expression analysis in articular cartilage and GP tissues

The articular cartilage and GP were laser micro-dissected (LMD7, Leica, Microsystems Inc., Buffalo Groove, IL) and purified using RNAeasy micro kit (Qiagen). The cDNAs were synthesized cDNA with the EcoDryTM Premix (Clontech, Mountain View, CA) in a 20µl reaction volume. Quantitative PCR (qPCR) was performed using TaqMan FastAdvanced Master Mix (Applied Biosystems, Foster City, CA). The probes were purchased from Integrated DNA Technologies, Inc. Coralville, IA) as follows; *Acan*: Mm.PT.58.10174685, *Col9a1*: Mm.PT.58.6363324, *Ihh*: Mm.PT.58.30489545, *Mmp13*: Mm.PT.58.42286812, *Tnfsf11*: Mm.PT.58.29202697 and *Actb*: Mm.PT.39a.22214843.g. Reactions were performed with 3 biological replicates. Changes in transcript abundance were calculated using the ΔΔCt method with *Actb* as the reference transcript following the manufacturer protocol.

### Statistical analysis and sample size

Results were analyzed by unpaired and non-parametric test or ordinary one-way ANOVA and Turkey’s multiple comparison test using Prism 10 (GraphPad Software, La Jolla, CA). Sample size was calculated with power 0.80, α=0.05 and the expected difference ranging 20-30%, and revised according to the data.

## Results

### Local application of RARγ agonist-loaded nanoparticles inhibits GP activity

In previous studies we showed that selective agonists for RARγ are able to reduce endochondral bone formation in mouse models of acquired and genetic heterotopic ossification.^20^ One drug, NRX204647, was particularly potent and was chosen here to generate unequivocal proof-of principle evidence for the soundness of our central thesis.

NRX-204647 was loaded into biodegradable polylactic acid (PLA)-NPs. The *in vitro* bio-assays revealed that the biological activity of NRX404647 was retained after entrapment within the PLA-NPs (Figure 1A). The NRX204647-loaded NPs (NRX-NPs) did continue to release active NRX204647 up to 192 h (8 days) in vitro at 37°C (Figure 1B), indicating that this formulation allows sustained drug release for at least a week.

**Figure 1.**
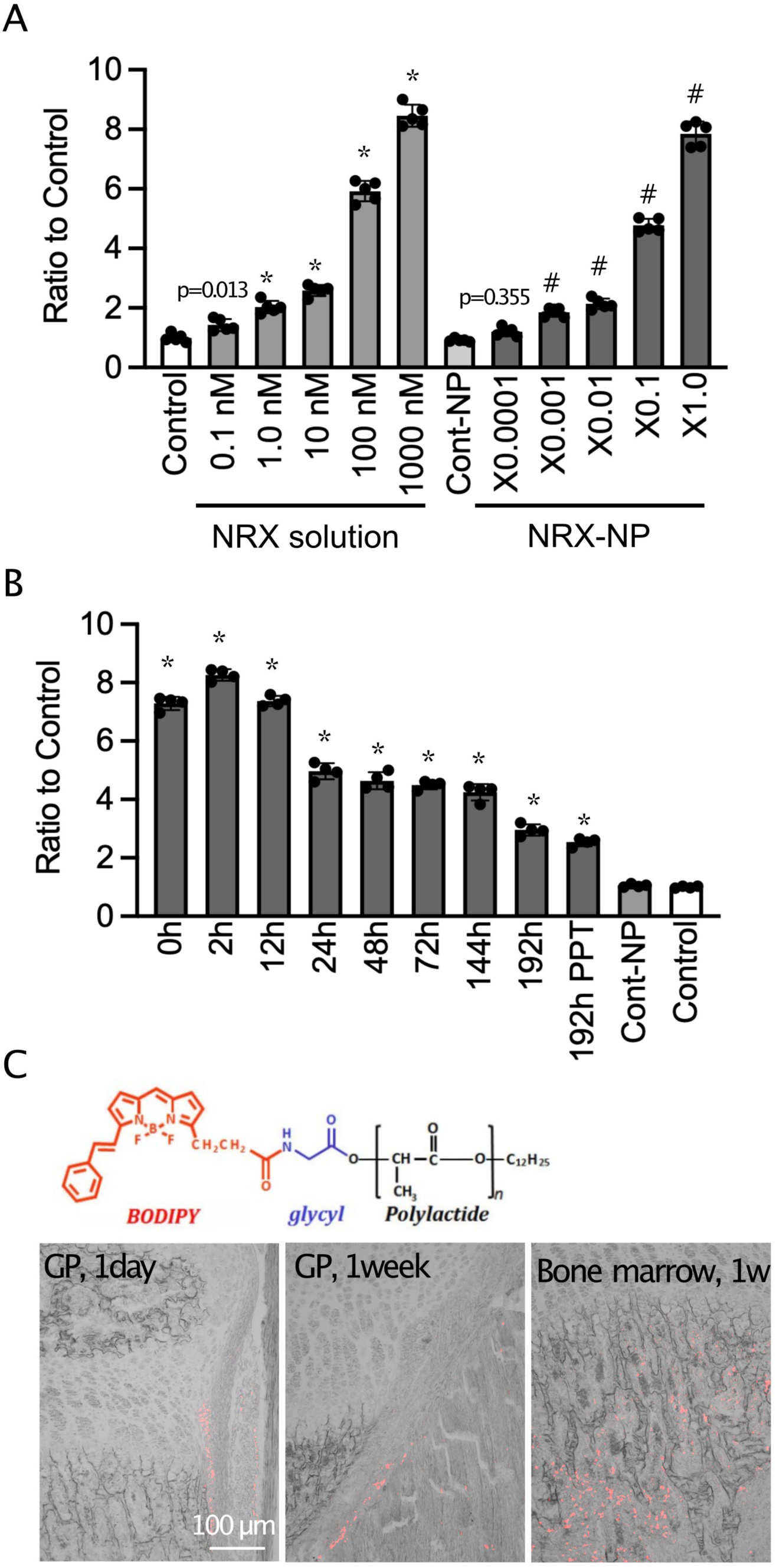
The nanoparticles sustain long release of NRX204647 and retention. A. The RARγ-reporter HEK293 cells were treated with a serially diluted NRX204647 (NRX solution) or NRX-NPs. The luciferase activity was measured 48 hrs after treatment using the luciferase assay kit. Individual values, average and standard deviation (SD) are shown. * p < 0.0001 to vehicle solution (Control), #p<0.0001 to control nanoparticles (Cont-NP). B, The NRX-NPs were incubated in the chondrocyte culture medium (high glucose DMEM containing 10% FBS) for 2 hrs at 37°C and collected by centrifugation at 100,000 rpm for 15 min at 4°C. The supernatant was stored at 4°C until use. The precipitated NRX-NPs were resuspended in the fresh culture medium and further incubated. This step was sequentially repeated to have NRX-NP supernatants after 12-, 24-, 48-, 72-, 144- and 192-hrs incubation. The RARγ-reporter HEK293 cells were treated with the initial NRX-NPs solution (0h), the NRX-NP-supernatants obtained after the indicated time incubation or the precipitated NRX-NPs (192h PPT) after 192-hrs incubation. The luciferase activity was measured 48 hrs after treatment using the luciferase assay kit. Individual values, average and standard deviation (SD) are shown. * p < 0.0001 to vehicle solution (Control). C, The BODIPY fluorescent-labeled NPs were injected into the vicinity of the proximal growth plate (GP) or bone marrow of the tibia. The tibias were harvested 1 day or 1 week after injection, and the distribution of the fluorescent-labeled NPs (red) were examined.

BODIPY-labeled NPs (Figure 1C) were implanted by injection in the vicinity of proximal GPs or into bone marrow for comparison. Fluorescence signal emanating from the BODIPY-labeled NPs was readily detectable at the injected proximal tibial site (Figure 1C, GP 1 day) and remained detectable locally for up to1 week from injection (Figure 1C, GP 1 week). To ensure that the drug-loaded NPs affected flanking tissues only, we injected tamoxifen-loaded NPs near the proximal GP on the medial side of left tibiae in *Ai6* reporter mice. Three days after injection, left and right tibiae were harvested and processed for fluorescence signal detection. Left tibia GP displayed large number of reporter-positive cells while the right tibia GP had barely discernable signal (Supplementary Figure S1), indicating that most, if not all, of tamoxifen released from NPs acted locally and on flanking tissues.

Next, we injected NRX-NPs to the proximal growth plate of left tibiae in juvenile 3 weeks-old C57BL/6j mice. The contralateral tibia in each mouse was implanted with blank NPs, serving as control of the procedure. NRX-NPs (0.8 μg/μl, 2ul/side) were implanted near the medial and lateral sides of proximal GPs on left tibiae (Figure 2A). The same volume of control NPs were injected near the GP of contralateral tibia. NRX-NP-implanted mice grew as much as control mice by 5 weeks of age (Figure 2B). However, radiographic imaging revealed that the GP in control mice was open and radiolucent (Figure 2C, Blank-NP. arrow) while the GP in NRX-NP-injected tibiae was not clearly visible (Figure 2C, NRX-NP, arrow). Tibiae were dissected for phenotypic assessment (Figure 2D). We found that the overall length of NRX-NP-injected tibiae was significantly shorter compared to that in control blank-NP injected mice (Figure 2E, NP).

**Figure 2.**
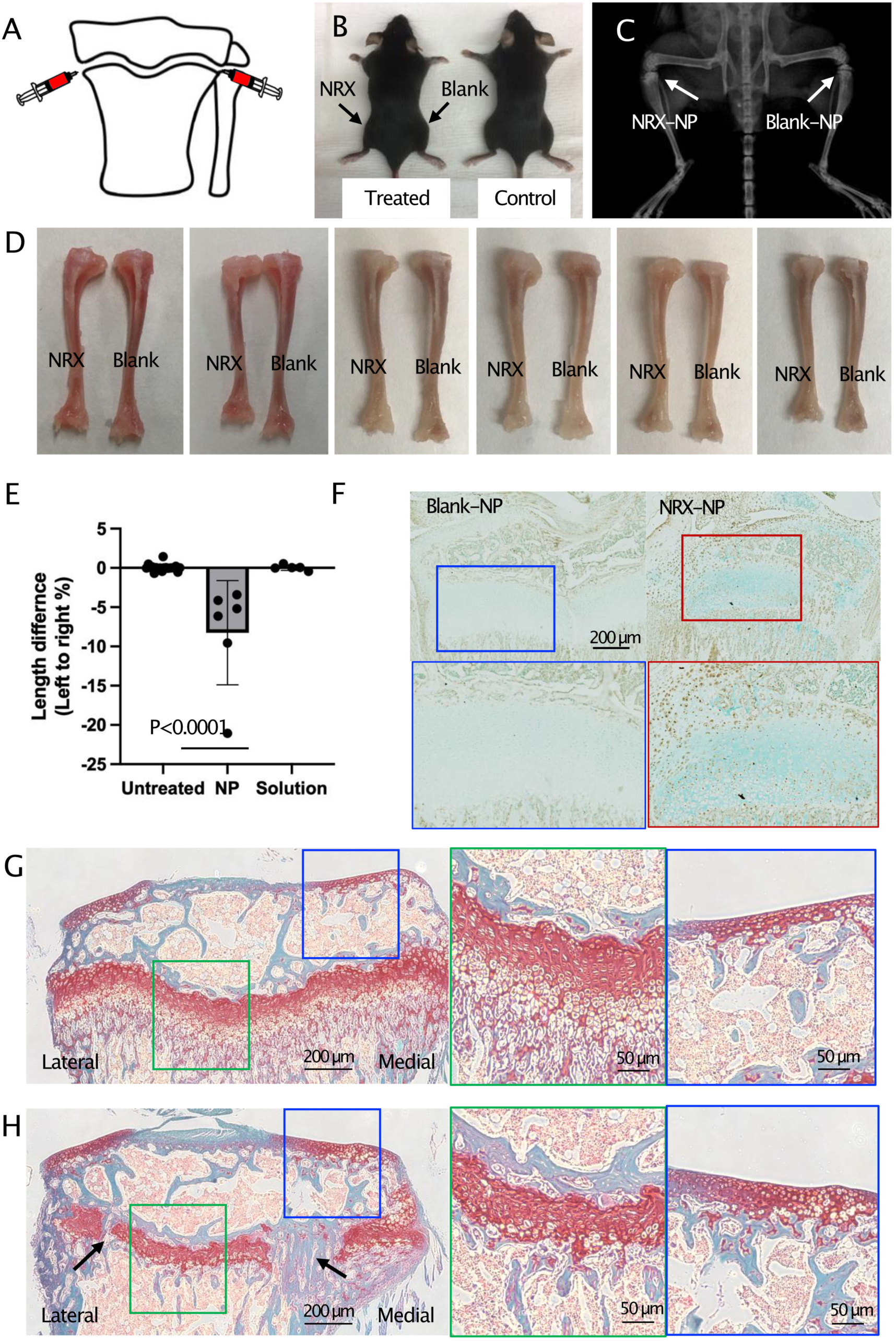
Local injections of the NRX-NPs induce the targeted growth plate closure. A, The NRX-NPs (NRX, 1.6 μg/2 μl/site, weekly) or unloaded-NPs (Blank) were injected at the medial and lateral sides in the vicinity of the proximal tibia growth plate in 3-weeks old C57BL/6J female mice (n=6). B, Gross appearance of the NP-treated or nontreated mice (5 weeks of age) 2 weeks after the treatment started. C, The representative radiographic image of the NP-treated mice (5 weeks of age). D-H, The tibiae treated with NRX-NPs (left) and Blank-NPs (right), or NRX204647 (left) and vehicle (right) solution were harvested at 5 weeks of age. (D) Gross appearance of the bones after removal of the soft tissues. (E) The bone length was measured using the radiographic images. The percentage ratio of left tibia length to right tibia length was calculated. Individual values, average and standard deviation (SD) are shown. (F) Tgm2 immunohistochemistry staining images of the Blank-NP- and NRX-NP-implanted growth plate 2 days after implantation of the NPs. (G and H) The representative images of the Blank-NP-(G) and NRX-NP (H)-implanted growth plate of the proximal tibiae. The frontal sections were stained with Safranin O.

Untreated left and right tibiae were of comparable length (Figure. 2E, Untreated). An equivalent amount of NRX202647 solution was injected in the same manner as for NRX-NPs, but the difference in length between tibiae injected with NRX solution and those injected with the control vehicle was marginal (Figure 2E, Solution). The NRX-NP-implanted GP showed strong immunoreactivity to the antibody against Tgm2, a direct target of retinoid signaling^33^ compared to the Blank-NP-implanted GP, indicating that growth plate chondrocytes respond to NRX202647 (Figure 2F). Histological examination revealed that the GPs treated with NRX-NPs were visibly affected. The Blank-NP implanted growth plate appeared normal while the NRX-NP implanted growth plate was shortened (Figures 2G vs 2H). In most cases of NRX-NP implanted growth plate, some regions displayed a complete closure (Figure 2H, arrows), and other regions had lost the hypertrophic zone and cartilage remnants in the primary spongiosa (Figure 2H, green square). The articular cartilage of the NRX-NP-implanted epiphysis showed immunoreactivity to Tgm2 (Figure 2F, left), but showed similar histology to that of the Blank-NP implanted articular cartilage (Figures 2G vs 2H, blue squares).

To determine whether the NRX-NP approach could elicit even finer morphogenic effects, we implanted them only on the lateral side of left tibia GPs (Figure 3A) and monitored responses over time. Importantly, we found that the epiphyseal end of tibiae became tilted laterally (Figure 3B). Average difference in medial proximal tibial angle (MPTA) was about 6.8° at the 2-week time point compared to control contralateral tibiae (Figure 3C). The lateral side of the NRX-NP implanted GP was closed whereas the medial side has shorter GP losing hypertrophic zone compared to the Blank-NP-implanted GP (Figure 3E), showing that implantation of NRX-NPs at the lateral site produced a gradient effect on the growth plate, but even the medial site of the growth plate was affected. The data indicate that unilateral application of the retinoid drug could enable clinically-relevant manipulation of epiphyseal end/joint angle alignment and re-alignment, but improved techniques for controlling drug localization are still needed.To delineate the possible consequences by the implanted NRX-NPs, we examined expression of major cartilage genes by quantitative PCR, using laser-captured samples of GP and articular cartilage from each mouse. NRX-NP-implanted GPs exhibited a major decrease in gene expression levels of *Acan*, *Col9a1* and *Ihh* and a concurrent increase in expression of *Mmp13* and *Tnfsf11*(*Rankl*) compared with those in the GPs in untreated control mice (Figure. 4A). In contrast, there were no measurable change in expression of *Acan*, *Col9a1*, *Ihh, Col10a1*and *Mmp1*3 in articular cartilage above the GPs implanted with NRX-NPs or control NPs (Figure 4B), attesting to the specific action of the NPs restricted to the site of implantation.

**Figure 3.**
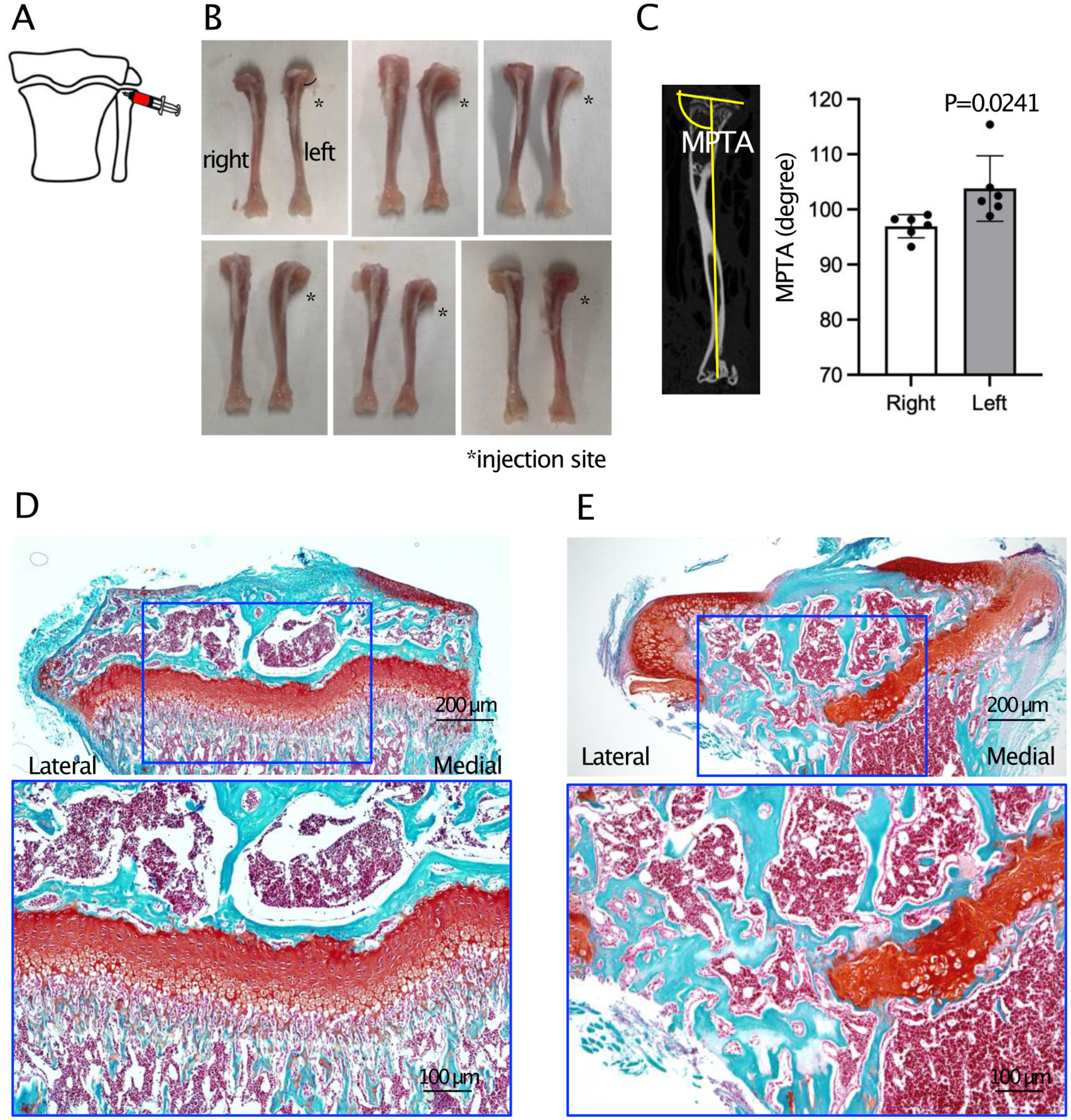
Unilateral injections of the NRX-NPs tilt the epiphyseal end of the targeted tibia. A, The NRX-NPs (NRX, 1.6 μg/2 μl/site, weekly) were injected at the lateral sides in the vicinity of the proximal growth plate of left tibiae in 3-weeks old C57BL/6J female mice (n=6). B, Gross appearance of the NP-treated (*left*) or contralateral nontreated (*right*) tibiae (5 weeks of age). C, The medial proximal tibial angle (MPTA), the medial angle between the tangent to the tibial plateau line and the mechanical axis of the tibia line was drawn and measured (yellow). Individual values, average and standard deviation (SD) are shown. D and E, The representative images of the Blank-NP-(D)-implanted or nontreated growth plate of the proximal tibiae. The frontal sections were stained with Safranin O.

**Figure 4.**
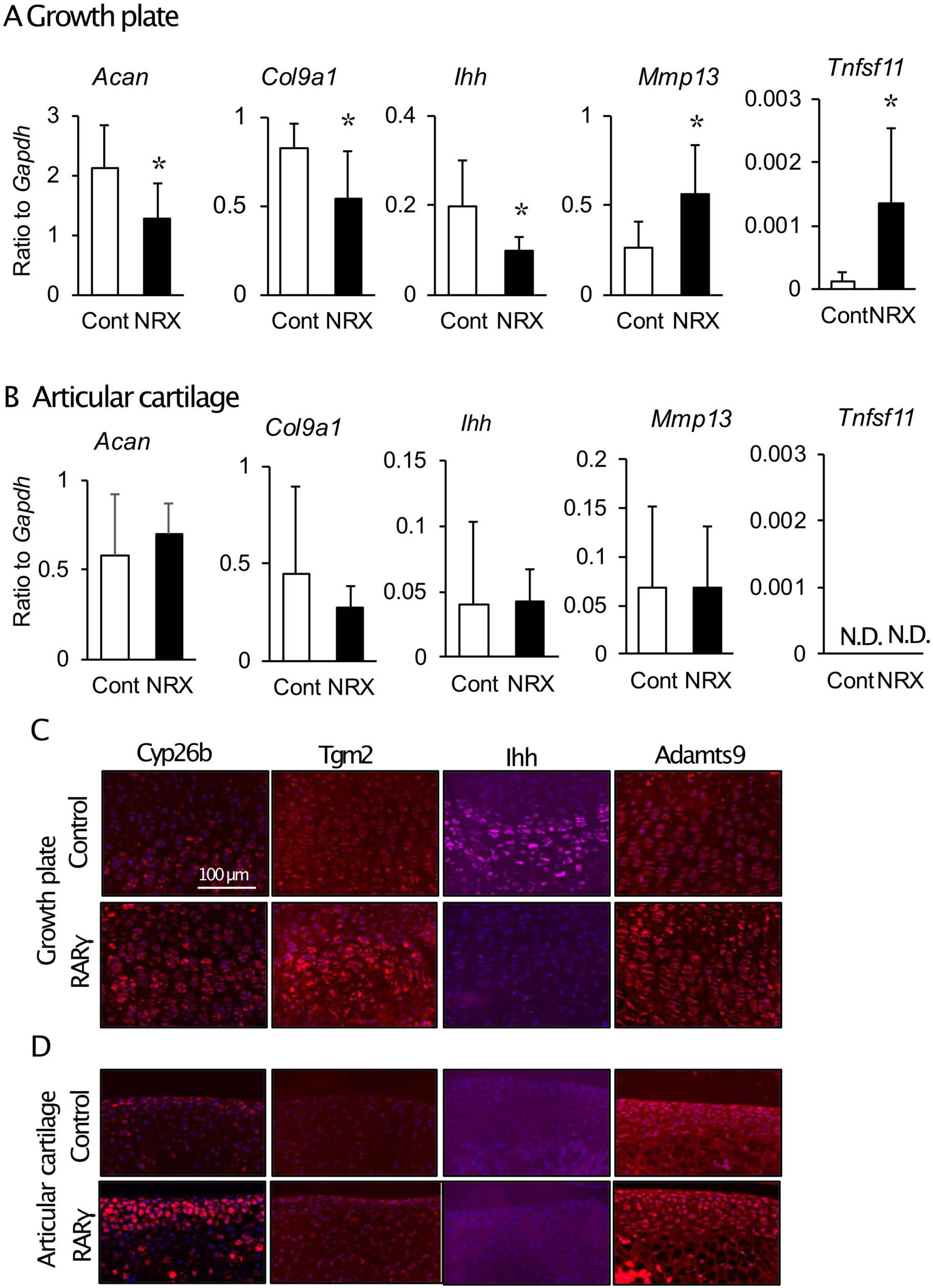
Differential responses to RARγ agonist treatment between growth plate and articular cartilage. The NRX-NPs (1.6 μg/2 μl/site, NRX) or unloaded-NPs (Control) were injected at the medial and lateral sides in the vicinity of the proximal tibia growth plate in 3-weeks old C57BL/6J female mice. The tibiae were harvested 4 days after the injection. A and B, The growth plate (A) and articular cartilage (B) were laser micro-dissected, and the total RNAs were prepared. Quantitative PCR (qPCR) was performed to examine gene expression of the indicated molecules. Reactions were performed with 5 biological replicates. Changes in transcript abundance were calculated using the ΔΔCt method with *Actb* as a reference transcript. Individual values, average and standard deviation (SD) are shown. Note no significant effects of NRX-NP treatment on articular cartilage. C and D, The growth plate (C) and articular cartilage (D) were subjected to *in situ* hybridization for indicated gene transcripts.

The changes in GP gene expression provoked by the RARγ agonist were further examined by in situ hybridization. Elevated transcript levels of typical RA target genes such as *Cyp26b* and *Tgm2* were detected in NRX-NPs-implanted GPs compared to control GPs (Figure 4C). Consistent with the *in vitro* results above, RARγ agonist treatment strongly reduced the amounts of *Ihh* transcripts whereas it greatly increased the amounts of *Adamts 9* transcripts (Figure 4C). These findings indicate that the changes seen in GPs following RARγ agonist treatment reflected a direct action by the drug on GP chondrocytes. Interestingly, an increase in *Cyp26b* transcripts was also detected in articular cartilage above the NRX-NPs-implanted GPs, but upregulation of *Tgm2 were marginal,* and *Adamts9* transcript levels had remained at control levels (Figure 4D,. The data indicate that RARγ agonist possibly diffusing away from the GP implantation site toward the epiphyseal end may up-regulated target molecules of retinoid signaling, such as Cyp26b and Tgm2 although histology was similar between control and RARγ-treated articular cartilage (Figures 4G vs 4H, green squares). It is also possible that the biological responses to RARγ agonist of articular cartilage and GP are different, with the latter being more sensitive overall.

### RARγ agonists strongly affect chondrocyte phenotypic expression compared to agonists for RARα or RARβ

To better understand the action and selectivity of RARγ agonists on chondrocyte phenotype and function, epiphyseal chondrocytes isolated from newborn mice were seeded in standard monolayer culture and exposed to specific agonists of RARα, RARβ or RARγ, using natural retinoic acid (RA) for comparison since it activates all RARs.^19^ Chondrocytes treated with RARα agonist (AGN195183, 300 nM), RARβ agonist (BMS453, 300 nM) or vehicle exhibited a typical polygonal or round cytoarchitecture and produced and accumulated an abundant, refractile and seemingly normal matrix around them (Figure 5A, Control, RARα and RARβ). In comparison, those exposed to RARγ agonist (NRX204647, 100 nM) or RA (100 nM) became flat and lacked a refractile matrix (Figure. 5Α, RA and RARγ). Gene expression analysis demonstrated that RARγ agonist treatment strongly reduced gene expression of *Sox 9*, *Acan*, *Col9a1* and *Ihh* but stimulated expression of *Col10a1*, *Mmp13* and *Tnfsf11* (*RANKL*) all of which are normally increased in hypertrophic chondrocytes in the GP and involve the transition to bone.^2^ The effects of RARα and/or RARβ agonists on gene expression of *Sox 9*, *Col9a1*, *Ihh* and *Col10a1* were also detected, but the induced changes were milder compared to those by RARγ (Figure 5B). Furthermore, RARγ agonist treatment up-regulated the gene expression of matrix catabolic proteases including *Adamts8* and *Mmp 2,3, 9, 10, 11, 12, 13* and *15* while concurrently down-regulating gene expression of *Timp1* and *Timp3* (Supplementary Figure S2, WT and HT). The up-regulation of *Adamts*5 and *8* and *Mmp 2, 3, 9, 12, 13, 14* and *15* and the down-regulation of *Timp1* and *Timp3* by the RARγ agonist were specific since such gene expression changes were not detected in genetically mutated *RARγ*-deficient chondrocytes (Supplementary Figure S2, ΚΟ).

**Figure 5.**
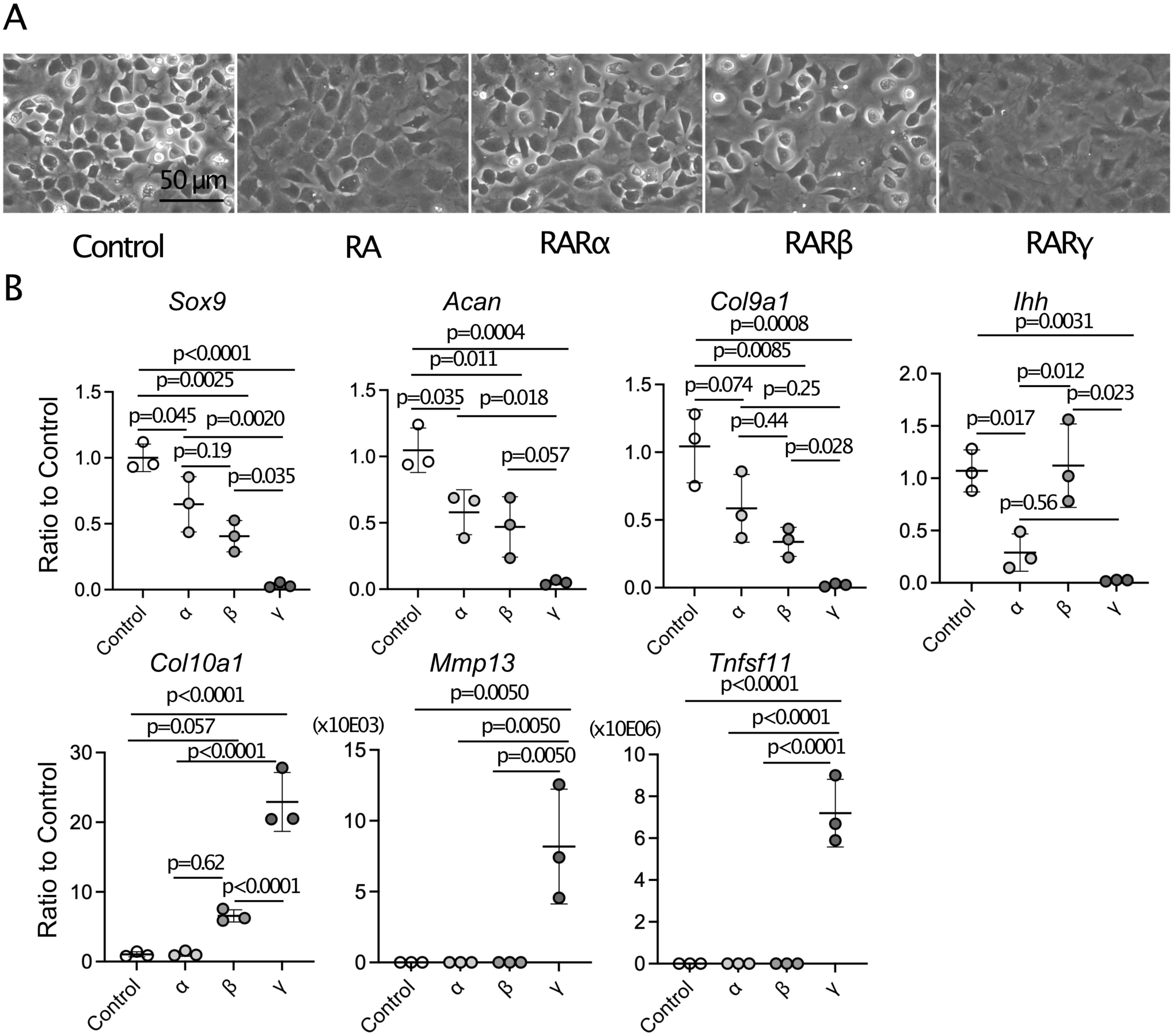
RARγ agonists strongly affect cultured chondrocyte function. Epiphyseal chondrocytes isolated from neonatal mice were treated with vehicle ethanol, retinoic acid (100 nM, RA), RARα agonist (AGN195183, 300 nM), RARβ agonist (BMS453, 300 nM) or RARγ agonist (NRX204647, 100 nM) 2 days after plating. A, Phase contrast images 4 days after treatment. B, Total RNAs were prepared 4 days after treatment and subjected to qPCR to examine expression levels of *Sox 9*, *Acan*, *Col9a1*, *Ihh*, C*ol10a1*, *Mmp3* and *Tnfs11* (*Rankl*). n=3/group. Individual values, average and standard deviation (SD) are shown.

### Systemic administration of RARγ agonists has broader skeletal consequences

To verify the selective effects of RARγ agonists on growth plate chondrocytes in vivo, as seen in the in vitro cultures described above, we set out the in vivo experiments to test the systemic effects of retinoid agonists and antagonists. Groups of mice were administered RARα agonist (AGN195183, 4 mg/kg), RARβ agonist (BMS453, 4 mg/Kg), RARγ agonist (NRX204647, 1 mg/Kg) or RARγ antagonists (CD2665 or 7C, 4 mg/kg) every other day from 11days old until harvest. Mice treated with RARγ agonist experience overall skeletal growth retardation by day 8 from the start of treatment whereas those receiving RARα or RARβ agonist did not (Figure 6A). Radiological images of hindlimbs revealed that the GPs were closed in RARγ agonist-treated mice (Figure 6B, RARγ, arrows) but appeared open and had radiolucent in mice receiving RARα agonist, RARβ agonist or vehicle control (Figure 6B, control, arrows). Body length was affected only in the RARγ agonist-treated group (Figure 6C, RAR agonists). The longitudinal growth of femur and tibia parameters equally affected in both males and females (Figure 6D). While RARγ agonists inhibited skeletal growth, administration of RARγ antagonists had minimal effects, trending toward a stimulation of growth (Figure 6C, RARγ antagonists).

**Figure 6.**
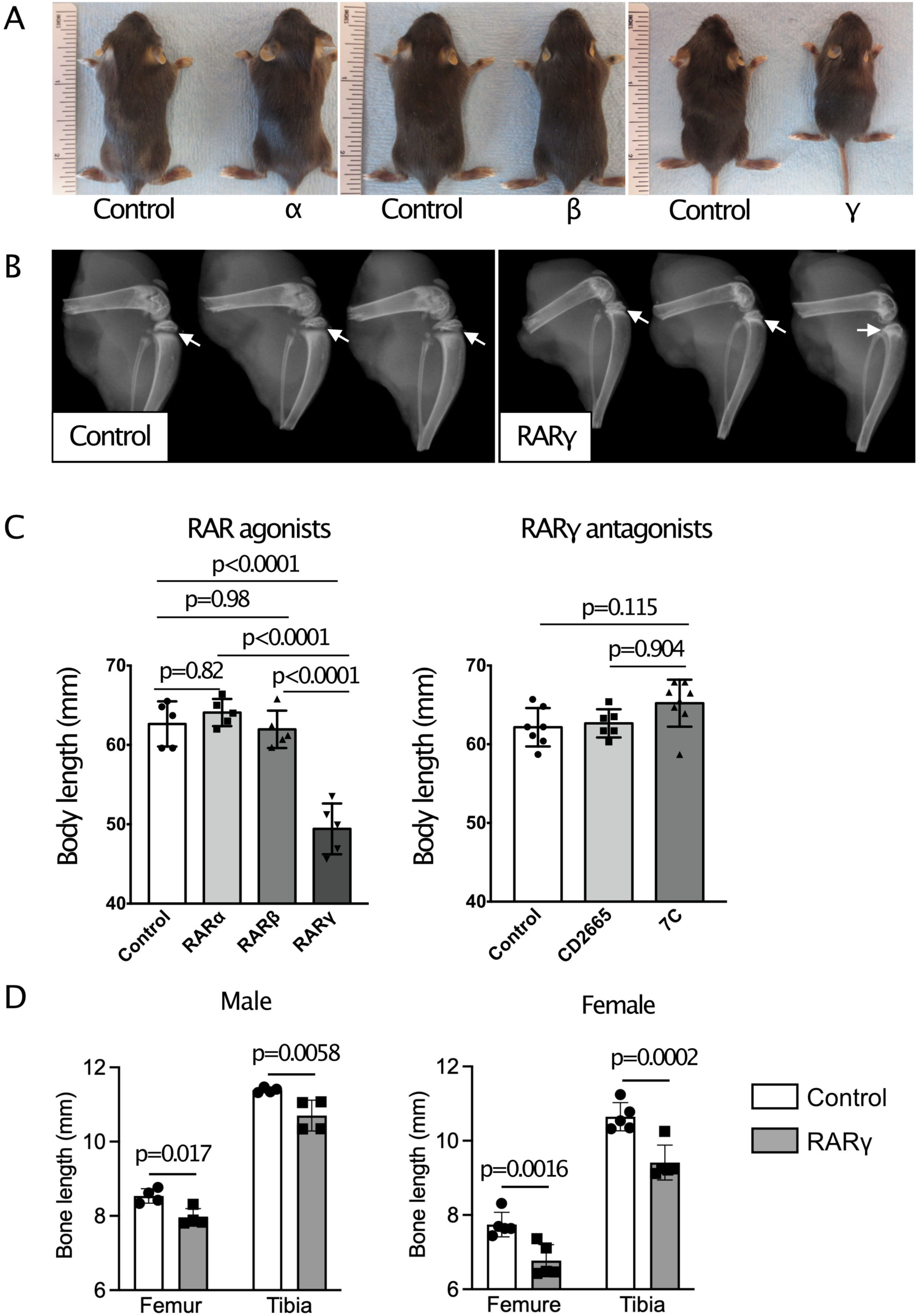
RARγ signaling dominantly affects skeletal growth. A-C, C57BL/6J female mice were treated with corn oil (Control), selective agonists of RARα (AGN195183, 4 mg/kg), RARβ (BMS453, 4 mg/Kg), RARγ (NRX204647, 1 mg/Kg) or antagonist for RARγ (CD2665 and 7C, 4 mg/kg) via peritoneal injections (11 days of age) or gavage (over 14 days of age) starting at 11 days of age every other day. The hind limbs were harvested at and subjected to radiological examination. (A) Gross appearance 8 days after initial treatment (19 days of age). (B) Radiological views of the knee joints 8 days after initial treatment (19 days of age). Arrows indicate the proximal tibial growth plate. Note RARγ-agonist treated group (right panel) showed closure of growth plate. (C) Body length was measured 8 days after initial treatment (19 days of age). D, C57BL/6J female and male mice were treated with corn oil (Control) or selective agonists of RARγ (NRX204647, 1 mg/Kg) via peritoneal injections starting at 14 days of age every other day. The hind limbs were harvested at 22 days of age and subjected to radiological examination. The tibia and femur lengths were measured by ImageJ. Individual values, average and standard deviation (SD) are shown.

Histological examination confirmed that GP height was appreciably reduced within 3 days of standard RARγ agonist treatment and that the GP was closed by day 8 (Supplementary Figure 3A). Furthermore, a strong reduction in cartilage matrix was detected by day 5 (Supplementary Figure 3A). The RARγ treatment also inhibited cell proliferation, increased the height of the hypertrophic zone and stimulated proteoglycan degradation as determined by EdU incorporation activity, immunostaining for collagen 10 and immunostaining for aggrecan neo-epitopes caused by MMP action, respectively (Supplementary Figure S3B-3D). These observations were consistent with the changes in gene expression found in the NRX-NP-implanted GP and NRX204647-treated chondrocyte cultures (Figures 4A and 4C).

## DISCUSSION

As we point out above, pediatric fractures involving the GP remain a major clinical challenge and require careful and complex surgical intervention and follow-up clinical visits. This is because such fractures can lead to either inhibition or acceleration of bone growth, resulting in progressive growth imbalances, skeletal deformities, decreased motion and/or chronic pain.^7,8^ When the difference in limb length is substantial the condition is referred to as limb length discrepancy (LLD) emphasizing its potential to cause severe problems with mobility and skeletal growth and functioning.^34^ Asymmetrical dysfunction of a GP in a single long bone can cause angular deformities such as in hindlimbs (genu valgum or genu varum). When LLD and angular deformities become severe or cause pain and/or gait disturbances, growth modulation is attempted by surgery such as epiphysiodesis performed by physeal ablation, transphyseal screws or staples, or reversible epiphysiodesis with a tension elastic band.^35^ If the related problems worse, an osteotomy may be performed to correct the deformity and imbalance. Currently, surgical interventions remain the only means of definitive treatment for these conditions.^9^ Surgery is invasive, gives substantial burden to the affected children and their families and may lead to complications.

This study was thus aimed to develop an alternative pharmacological method of intervention, and we specifically tested whether local application of RARγ agonist would restrain GP function, possibly eliciting an impact on the GP similar to that of a surgical intervention such as epiphysiodesis. To exert local action and limit side effects, we have developed a NP-based drug delivery approach. This has been explored for treating musculoskeletal disorders such as osteoporosis,^36^ rheumatoid arthritis^37,38^ and osteoarthritis.^39,40^ NPs formulated with bioactive agents have also been studied in combination with scaffolds on skeletal tissue defects to induce bone and cartilage regeneration.^41,42^ Our *in vitro* data demonstrate that the NRX-NPs are able to steadily release active retinoid drug for at least 192 hrs (Figure 1), indicating that the PLA-based NP is an effective and appropriate drug cargo system for local and sustained delivery of the synthetic retinoid. The data are in line with our *in vivo* observations demonstrating that local application of NRX-NPs leads to GP closure and restricts the lengthening of the targeted long bone while the contralateral element remains normal and unaffected (Figure 2). These effects are most likely due to direct drug action on the GP chondrocytes themselves rather than a secondary consequence of local drug action. This is sustained by our *in vivo* and *in vitro* data showing that NRX204647 is able to readily modulate the phenotype of GP chondrocytes, evident by the decreases in matrix gene expression and the increases in gene expression of hypertrophic chondrocyte traits (Figures 4 and 5B). Support comes also from our findings that RARγ agonist-treated GPs up-regulated the expression of retinoid target genes *Cyp26b1* and *Tgm2* (Figure 4C). Because the effects elicited by RARα or RARβ agonists were in general much milder, the data sustain the important notion that RARγ is the critical RAR family member operating in GP chondrocytes.^14^

Especially, we show here that when implanted in one side of tibia GP only, the NRX-NPs were able to change the angulation of the adjacent epiphyseal end and joint (Figure 3). These results provide additional proof-of-principle evidence that the NP-based drug-delivery approach can be used for even finer and more restricted manipulation of key morphological features of the long bones such as angulation. Limitations of experiments with local injections of NRX-NPs include the need for detailed spatial and temporal analysis of histological and functional changes in the affected growth plate and the need to determine sex and tissue differences in sensitivity to NRX-NPs. Nevertheless, it appears that our retinoid local delivery technique could aid or even possibly replace corrective surgeries and be useful to improve relatively mild LLD cases and angular deformities that are currently not considered surgical cases. A pharmacologic intervention would reduce the patient’s future risk to develop joint problems over time such as osteoarthritis.^7–9^ An additional advantage would be that the drug therapy could be used to fine-tune growth inhibition by adjusting drug concentration and/or frequency depending on LLS severity and individual susceptibility to the drug although further research may be required using larger animals and advanced drug delivery tools.

It is important to emphasize our observations that systemic administration of RARγ antagonist treatment did not significantly impact the GP and skeletal growth under the regimens used. In a previous study, we showed that RARγ antagonists can actually rescue GP closure caused by systemic treatment with hedgehog inhibitory drugs used in cancer therapies.^26^ Thus, it should be studied if NPs filled with a RARγ antagonist be used to counteract conditions involving pathological impairment of GP activity and function. Together, the above data and considerations raise the novel possibility that implantation of NPs impregnated with a selective RARγ antagonist or agonist could be used in a variety of clinical settings aiming to correct long bone length, physical alignment and/or angulation by acting with high site-specificity on the neighboring GPs. Palovarotene, other RARγ agonist, was recently approved by Health Canada and U.S. Food and Drug Administration (NDA215559) for the treatment of patients with Fibrodysplasia Ossificans Progressiva (FOP), genetic heterotopic ossification diseases and is given orally. The successful development of Palovarotene can be added to the ever expanding spectrum of natural and synthetic retinoid drugs in use or in testing for various pathologies,^43^ providing impetus to the current study aiming to establish a synthetic RARγ agonist such as NRX204647 as a promising local treatment to rectify GP defects.

The power and specificity of the NP approach are made even more apparent by our results following systemic drug administration. We observed a generalized skeletal growth retardation when the mice were given NRX-204647 systemically. This was actually predictable based on the fact that this drug is extremely potent^20^ and can elicit widespread effects if not contained and delimited locally. Skeletal growth retardation associated with ectopic cartilage formation was also observed when the clinically relevant RARγ agonist Palovarotene was given from 14 days old to mouse models of FOP.^44^ The systemic administration started from 11 days old in this study, but we did not find severe skeletal deformity (Figure 6B). This discrepancy may be explained by the different specificities of palovarotene and NRX204647 for RARγ, as well as the different routes of administration: NRX204647 has a higher affinity to RARγ than that of palovarotene, and NRX2024647 drug was administrated by gavage, except for a peritoneal injection on 11 and 13 days old while palovarotene was peritoneally injected throughout the treatment period in the study that reported skeletal deformity.^44^ Gavage administration is known to result in a slow and partial uptake of the drug and release into the circulation over a 24-hour period, whereas injections result in full and immediate drug availability and a burst of high drug levels and distribution in the circulation, eliciting acute and powerful but possibly deleterious effects.^45^ Notably, growth retardation by Palovarotene was not observed when it was given by gavage from older ages on the FOP disease model ^21^ and on the osteochondroma model ^46^. The growth plate activity deceases even during the growing stage,^47^ and the sensitivity of the growth plate to drugs may also decrease. Thus, patient age should be a critical factor for designing a drug regimen for RARγ agonists.

## Data availability

All data will be made available from the corresponding author M.I. upon reasonable request.

## Ethics approval statement

All animal experiments were approved by the Institutional Animal Care and Use Committees at Children’s Hospital of Philadelphia (959) and University of Maryland, Baltimore (0317003 and 0120005).

Translational study.

## Contributions

MI designed the experiments. MM, KU, IA, YU, TO, ML and MI conducted experiments; MM, SO, JMA, JH, MP, MEI and MC evaluated the results; and MM, MI and MEI prepared the manuscript with revisions by JMA and MP. All authors have read and approved the final submitted manuscript.

## Funds

The work is supported by NIH grants AR056837 (MI/MP) and AR072713 (MI/MC), POSNA Angela S.M. Kuo Memorial Award, 823038 (JMA) and institutional funds.

## Acknowledgement

This study is supported by NIH grant R01AR056837 (MI) and R01AR072713 (MI and MC), POSNA Angela S.M. Kuo Memorial Award, 823038 (JMA) and the departmental fund (University of Maryland). We thank Dr. J. Mort (Shriners Hospital, Canada) for providing the VDIPEN antibody. MEI is a spouse of MI.

---

**Supplementary Figure S1.** Induction of Cre-recombination by local injection of 4-hydroxy tamoxifen (4-OH-tamoxifen) NPs. The incision was made to expose the left proximal epiphysis of the left tibia of 3-week-old Ai6 mice. 4-OH-tamoxifen-loaded nanoparticles (10 μg/2μl/stie) were injected at the medial side in the vicinity of the growth plate of the left proximal tibiae followed by suture of the incision. The injected tibia and contralateral tibia were harvested 3 days after the injection. The distribution of reporter proteins (ZsGreen) was evaluated in sagittal frozen sections. The fluorescent images of the medial and lateral sides of the growth plate were captured and superimposed with DAPI staining images.

**Supplementary Figure S2.** Analysis of gene expression of the extracellular matrix-related molecules changed by RARγ agonist treatment in cultured chondrocytes. Epiphyseal chondrocytes were isolated from homozygous (KO), heterozygous (HT) or wild type (WT) mice produced by mating of the heterozygous RARγ null mice. The freshly isolated cells were plated at the density of 1.2 x 10^5^/cm^2^ and reverse transfected on the PCR array plate (RT² Profiler™ PCR Array Mouse Extracellular Matrix & Adhesion Molecules, Qiagen). On the next day, the cultures were treated with RARγ agonists (NRX204647, 100 nM) for 48 hrs and subjected to PCR analysis following the manufacturer’s protocol. Average threshold cycle value (Ct value) was calculated from 4 wells and normalized to that of housekeeping gene *Gapdh*. The experiments were repeated twice independently.

**Supplementary Figure S3.** RARγ agonist treatment induce closure of growth plate. C57BL/6j female mice were treated with corn oil (Control) and RARγ (NRX204647, 1 mg/Kg) via peritoneal injections (11 days of age) or gavage (over 14 days of age) starting at 11 days of age every other day. The tibiae were harvested 1, 3, 5 or 8 days after the initial treatment and subjected to histological inspection. A, Safranin O staining of the proximal tibial growth plate. B, EdU was injected 2 h in prior to euthanization 5 days after the initial treatment. The percentage of EdU-positive cells to total DAPI-positive cells was determined. C, Representative images of the immunostaining results for collagen 10 in the proximal tibia growth plate 5 days after the initial treatment. The collagen 10 fluorescence image is superimposed with DAPI and phase contrast images. D, Representative image of the immunostaining results for the neoepitope of cleaved aggrecan (VDIPEN) in the proximal tibia growth plate 5 days after the initial treatment. Counterstained with fast green.

