## Supplemental materials for "Retinoid-impregnated nanoparticles enable control of bone growth by site-specific modulation of endochondral ossification in mice"

Supplementary Figure S1 *Matsuoka et al.*

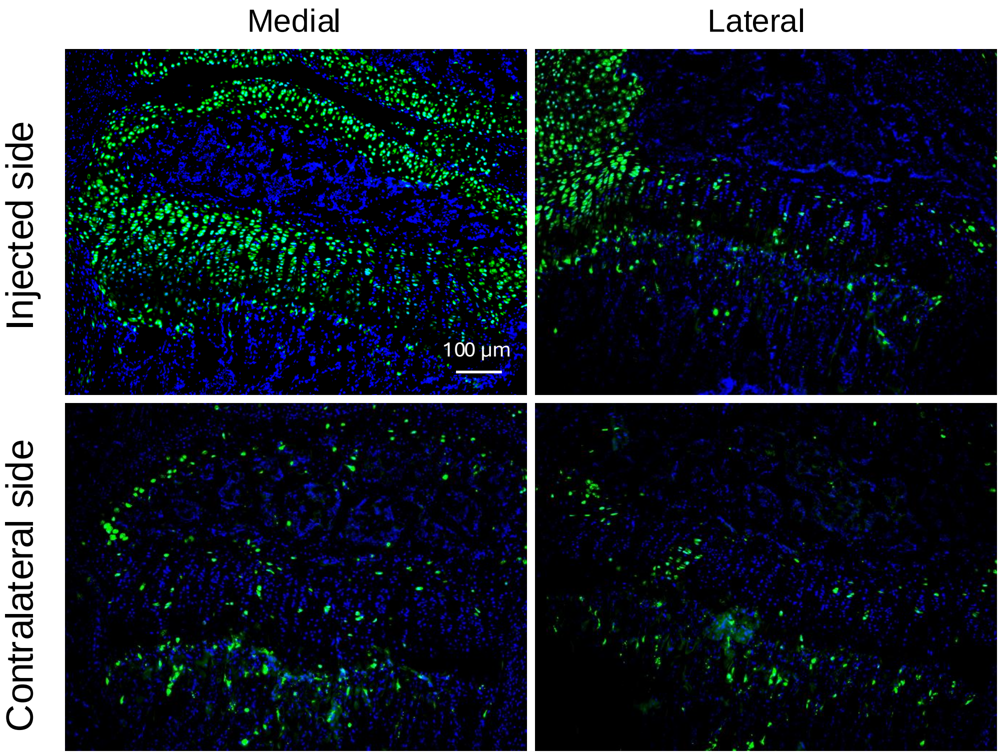


**Supplementary Figure S1.** Induction of Cre-recombination by local injection of 4-hydroxy tamoxifen (4-OH-tamoxifen) NPs. The incision was made to expose the left proximal epiphysis of the left tibia of 3-week-old Ai6 mice. 4-OH-tamoxifen-loaded nanoparticles (10 µg/2µl/stie) were injected at the medial side in the vicinity of the growth plate of the left proximal tibiae followed by suture of the incision. The injected tibia and contralateral tibia were harvested 3 days after the injection. The distribution of reporter proteins (ZsGreen) was evaluated in sagittal frozen sections. The fluorescent images of the medial and lateral sides of the growth plate were captured and superimposed with DAPI staining images.

Supplementary Figure S2. Matsuoka *et al.*

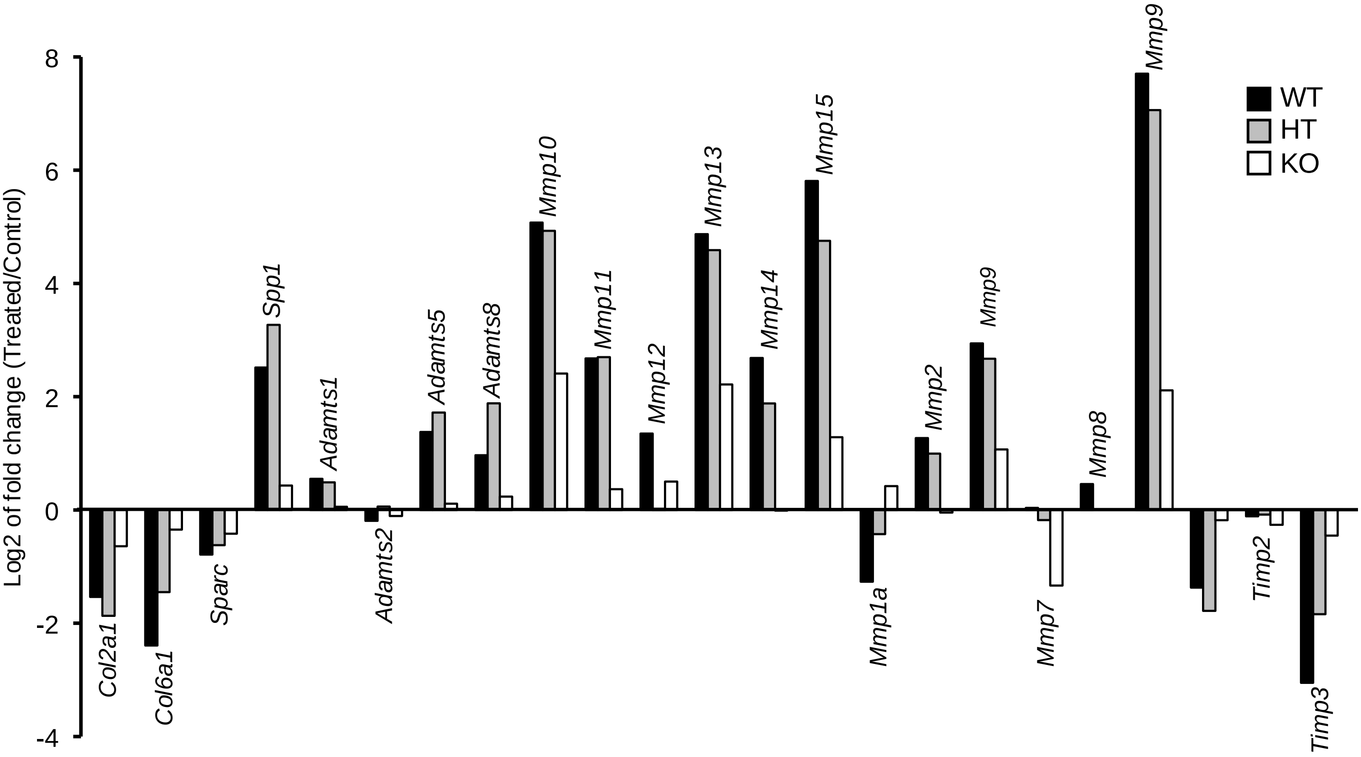


**Supplementary Figure S2.** Analysis of gene expression of the extracellular matrix-related molecules changed by RARγ agonist treatment in cultured chondrocytes. Epiphyseal chondrocytes were isolated from homozygous (KO), heterozygous (HT) or wild type (WT) mice produced by mating of the heterozygous RARγ null mice. The freshly isolated cells were plated at the density of 1.2 x $10^5$/cm$^2$ and reverse transfected on the PCR array plate (RT$^2$ Profiler™ PCR Array Mouse Extracellular Matrix & Adhesion Molecules, Qiagen). On the next day, the cultures were treated with RARγ agonists (NRX204647, 100 nM) for 48 hrs and subjected to PCR analysis following the manufacturer's protocol. Average threshold cycle value (Ct value) was calculated from 4 wells and normalized to that of housekeeping gene *Gapdh*. The experiments were repeated twice independently.

Supplementary Figure S3 *Matsuoka et al.*

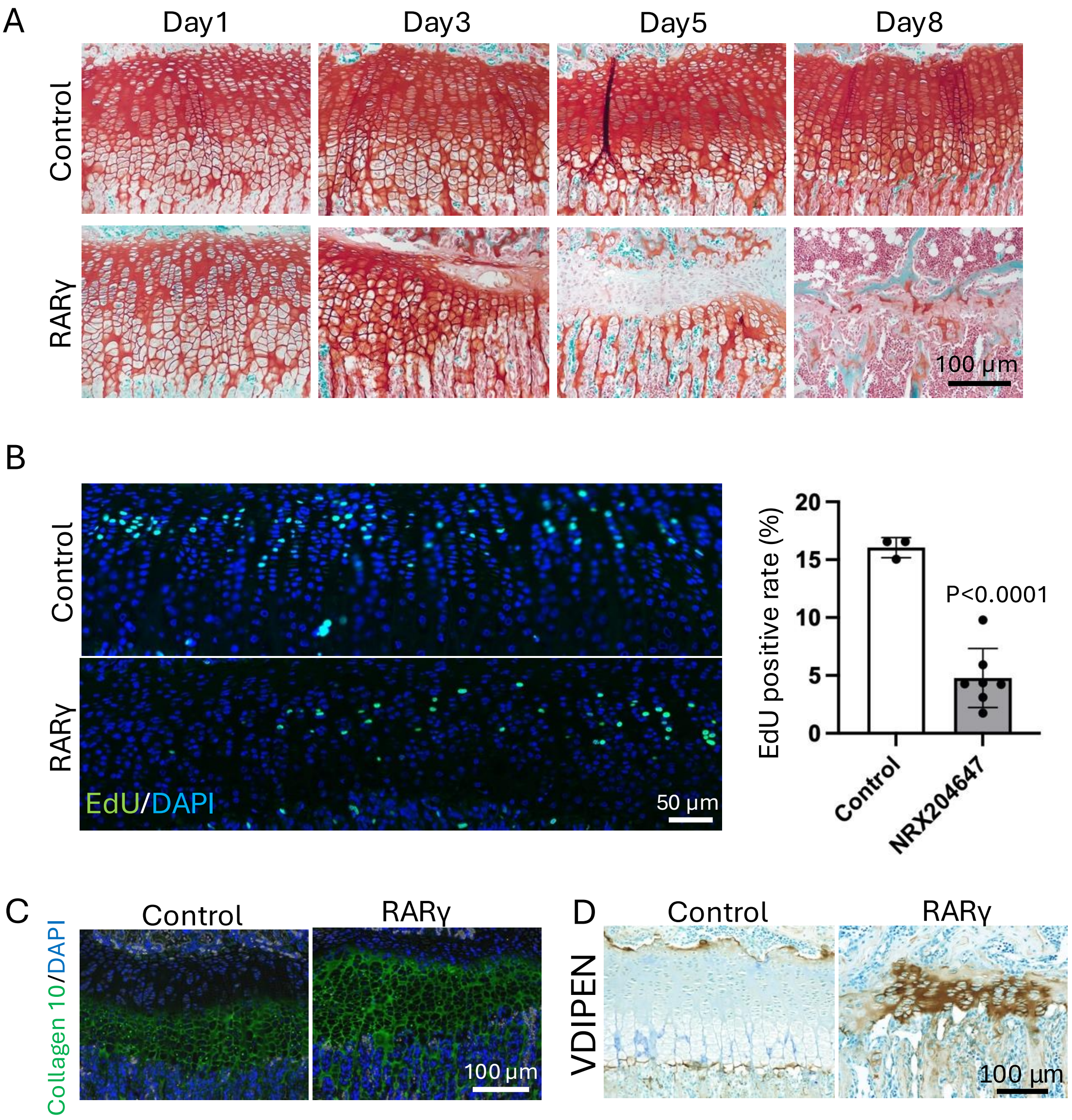


**Supplementary Figure S3**. RARγ agonist treatment induce closure of growth plate. C57BL/6j female mice were treated with corn oil (Control) and RARγ (NRX204647, 1 mg/Kg) via peritoneal injections (11 days of age) or gavage (over 14 days of age) starting at 11 days of age every other day. The tibiae were harvested 1, 3, 5 or 8 days after the initial treatment and subjected to histological inspection. A, Safranin O staining of the proximal tibial growth plate. B, EdU was injected 2 h in prior to euthanization 5 days after the initial treatment. The percentage of EdU-positive cells to total DAPI-positive cells was determined. C, Representative images of the immunostaining results for collagen 10 in the proximal tibia growth plate 5 days after the initial treatment. The collagen 10 fluorescence image is superimposed with DAPI and phase contrast images. D, Representative image of the immunostaining results for the neoepitope of cleaved aggrecan (VDIPEN) in the proximal tibia growth plate 5 days after the initial treatment. Counterstained with fast green.
